# Social experience alters different types of learning abilities controlled by distinct brain nuclei in *Kryptolebias marmoratus*

**DOI:** 10.1101/2021.04.25.441338

**Authors:** Cheng-Yu Li, Dietmar Kültz, Audrey K. Ward, Ryan L. Earley

## Abstract

Fighting experiences strongly influence aggressive behavior and physiology (winner-loser effects). These effects are conserved from invertebrates to vertebrates, but the underlying mechanisms remain unclear. Recent studies indicate that the brain social decision-making network (SDN) plays a key role in guiding experience-induced behavioral change. Also, while most studies have focused on how winning and losing experiences alter aggression, growing evidence points to these experiences driving multiple behavioral effects, including changes in the ability to learn. In mangrove rivulus fish (*Kryptolebias marmoratus*), we discovered that single winning experiences significantly improved spatial learning but not risk-avoidance learning, whereas single losing experiences drove the exact opposite to occur. These results provide strong evidence that winning and losing modulate diverse behaviors served by key nodes within the SDN, specifically the dorsolateral pallium (Dl; fish homolog to mammalian hippocampus, which serves spatial learning) and dorsomedial pallium (Dm; fish homolog to mammalian basolateral amygdala, which responds to fear). We therefore quantified whole-proteome expression within the forebrain (where Dm and Dl are located) of adult rivulus with divergent social experiences. We discovered 23 proteins were significantly differentially expressed in the forebrains of winners and losers. Differentially expressed proteins in losers related to modulation of cellular processes, apoptosis and learning while those in winners related to neuronal plasticity, neuroendocrine homeostasis, energy utilization, and learning. These results imply that winner-loser effects might be governed by very different patterns of protein expression, which could explain why winners and losers show such pronounced differences in behavioral performance.

**Significance Statement:** Social interactions permeate the daily lives of most animals and often result in changes in behavior for all parties. This implies that social experiences reorganize the brain in ways that promote the expression of alternative behaviors, or that help individuals cope with the outcome of such interactions. But how do aggressive interactions sculpt the brain at the molecular level? We used an emerging model organism, *Kryptolebias marmoratus*, to examine whether experiences modulate learning ability and then probe the potential neural mechanisms underlying these behavioral changes. We discovered that single winning and losing experiences dramatically altered spatial learning and risk-avoidance learning, respectively, indicating that winning and losing experiences have markedly different effects on the brain and cognitive processes.

**Classification:** Biological Sciences, Ecology

## Introduction

Engaging in aggressive interactions can potently alter physiology and future behavior. Social victory significantly increases aggression (1) and levels of androgenic hormones, like testosterone, in most taxonomic groups, including humans (2-5). On the other hand, social defeat dramatically decreases aggression (1) and can have chronic detrimental effects via prolonged activation of stress-related pathways in brain (6), which can induce depression and a variety of other behavioral disorders (7, 8). These responses to prior fighting experiences are termed the ‘winner effect’ and ‘loser effect’, respectively. Although winner-loser effects are strongly conserved from insects to humans, their underlying neurophysiological mechanisms remain largely unknown. Previous studies, which focus mainly on circulating hormones, indicate that testosterone and its receptor mediate aggression and winner effects (2, 4, 9-12), but recent studies suggest that other types of neural mechanism, such as activation of dopaminergic system (13, 14) within social decision-making network (SDN), might also govern behavioral responses to winning (15-17).

In addition, while most studies on contest behavior have focused on how winning and losing alter aggression, growing evidence points to these types of experience having manifold behavioral effects, including changes in cognitive ability (18-20). This suggests that social experiences might elicit widespread alterations to brain function. We thus took the approach of probing the activity of specific brain nuclei by examining the performance of animals on behavioral tasks known to be associated with activation of these brain regions. For instance, spatial learning ability has been associated with prior winning experience (19), which led to the investigating of hippocampus region, whereas the risk-avoidance learning ability has been associated with prior losing experience (18, 20), which led to scrutinizing basolateral amygdala. A recent study in fruit flies, *Drosophila melanogaster*, showed that social defeats induce a long-lasting loser effect associated with *de novo* protein synthesis (21), which is required for the formation and persistence of long-term memory, whereas social victories had no such long-term behavioral consequences and were not associated with *de novo* protein synthesis (21). These studies suggest that distinct neurobiological mechanisms might govern loser and winner effects across taxa. However, we know little about the memory-related neurophysiological mechanisms underlying behavioral responses to social competition (4, 22, 23) or the memory mechanisms that drive persistent changes in behavior following aggressive interactions (24, 25, 26). Therefore, our first aim was to explore temporal changes in aggression, spatial learning, and risk-avoidance learning in an emerging model organism, mangrove rivulus fish (*Kryptolebias marmoratus*, hereafter ‘rivulus’), following winning and losing experiences. We hypothesized that winners would show increased aggression and proficiency in spatial learning, because selection should favor behavioral processes that facilitate acquisition and defense of a territory and resources, which may require superior learning capacities in this context. We also hypothesized that losers would show decreased aggression but increased proficiency in risk-avoidance learning because selection should prioritize behaviors that facilitate recovery from aggressive contests, which may require avoiding risk and being less active overall.

Traditional social defeat paradigms (7), which expose animals to repeated losing experiences, are useful for identifying the terminal influences of multiple aggressive interactions on behavior and physiology. However, because the brain is exceptionally plastic, it is possible that the neural mechanisms responsible for short-term changes in behavior are different from those that maintain behavioral states over the long-term. While much attention has been given to exploring variation in neurobiology and behavior between animals occupying stable social rankings, there is increased awareness that examining responses to a single win or loss can provide insights into how the brain is initially reorganized by social inputs, and how and whether revamped neural (and associated behavioral) states are maintained over the long-term (27, 28). Because fighting experiences likely influence social behavior through a complex array of neuroendocrine interactions, investigating single candidate molecules is not sufficient to understand the mechanisms driving behavioral change. For this reason, and because proteins are ultimately responsible for producing the behavioral phenotype (29, 30), the second aim of this study was to quantify protein abundance in rivulus’ forebrain, which includes several brain nuclei (e.g., Dl - fish homolog of mammalian hippocampus; Dm - fish homolog of mammalian basolateral amygdala) implicated in modulating both aggression and learning, as well as physiological responses to acute social experiences. We hypothesized that winning and losing experiences would result in divergent, perhaps unique, patterns of forebrain proteome expression.

## Results

To investigate whether, and for how long, social experiences affect aggression and learning, we conducted a full factorial experiment with 3 experience and 3 decay-time levels. Fish (*n* = 675) were allocated to three experience treatments (W: winner; L: loser; N: no fighting experience [control]) and individuals in each experience treatment were subdivided into those exposed to the aggression, spatial learning or risk-avoidance learning tests (**Fig. 1A-C, *SI Appendix*, Movie S1-S5**). Animals were subjected to behavioral tests before (on Day 1; pre-experience behaviors) and 1h, 3h or 48h after social experience (on Day 15-17, based on pre-assigned decay-times; post-experience behaviors). Forebrain proteome expression was quantified in 12 additional individuals 1h after winning (*n* = 4), losing (*n* = 4) or control (*n* = 4) experiences, the time point at which fish showed pronounced changes in aggression and learning.

**Figure 1.**
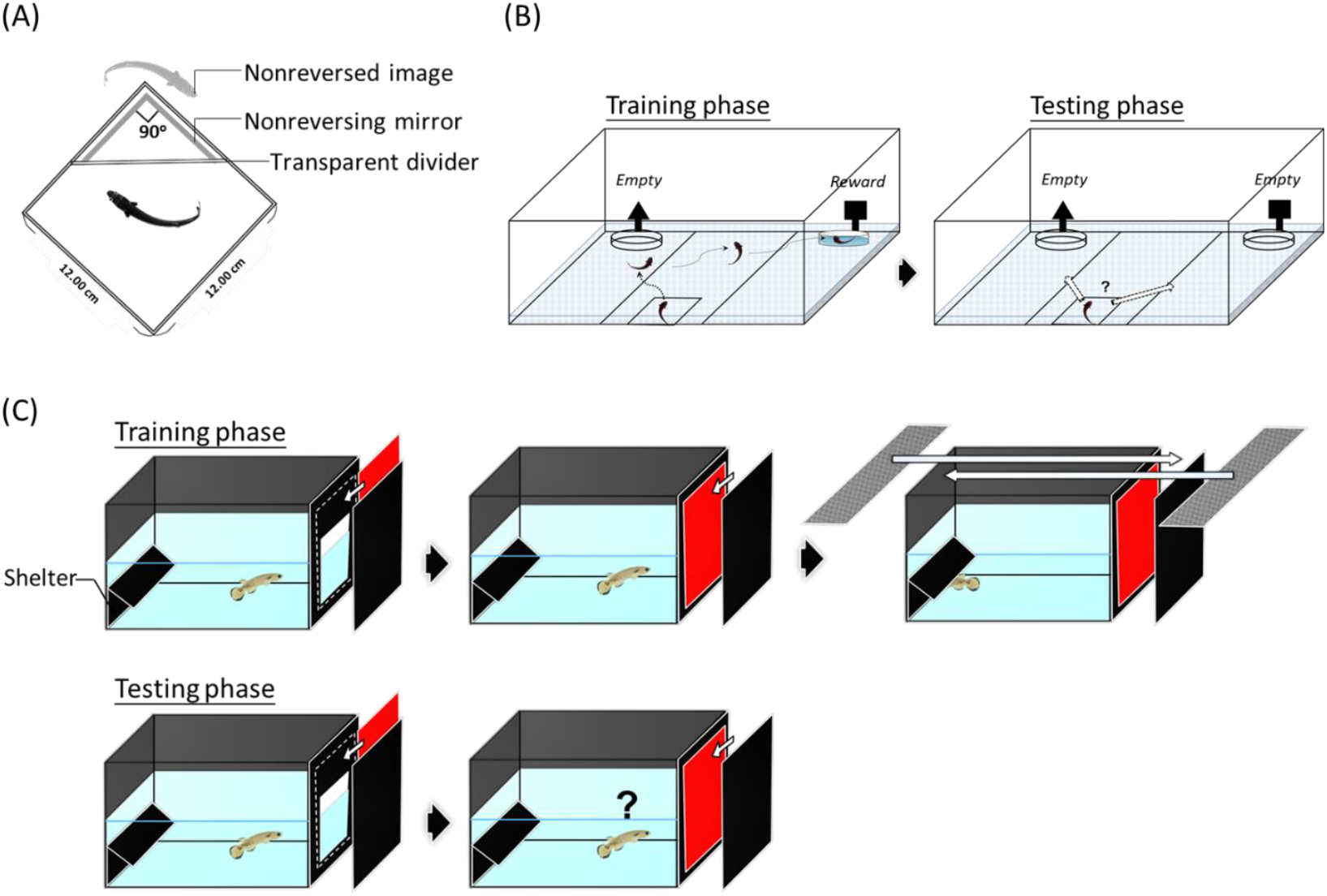
Setups for the **(A)** aggression test (non-reversing mirror image stimulation) **(B)** spatial learning test, which challenged fish to navigate an arena to find a reward (water/food) **(C)** risk-avoidance learning test, which challenged fish to associate red color with risk.

### Social experience and aggression

Before social experiences were obtained, individuals assigned to the 3 different experience treatments showed similar levels of aggression (**Fig. 2A, 2E, *SI Appendix*, Table S1**). However, different social experiences caused significant behavioral divergence (***SI Appendix*, Table S1**). Winners delivered more attacks to their mirror image than controls (*P* < 0.001, **Fig. 2F**), and losers were slower to launch first attacks (*P* < 0.001, **Fig. 2B**) and delivered fewer attacks to the mirror image (*P* < 0.001, **Fig. 2F**) than both winners and controls.

**Figure 2.**
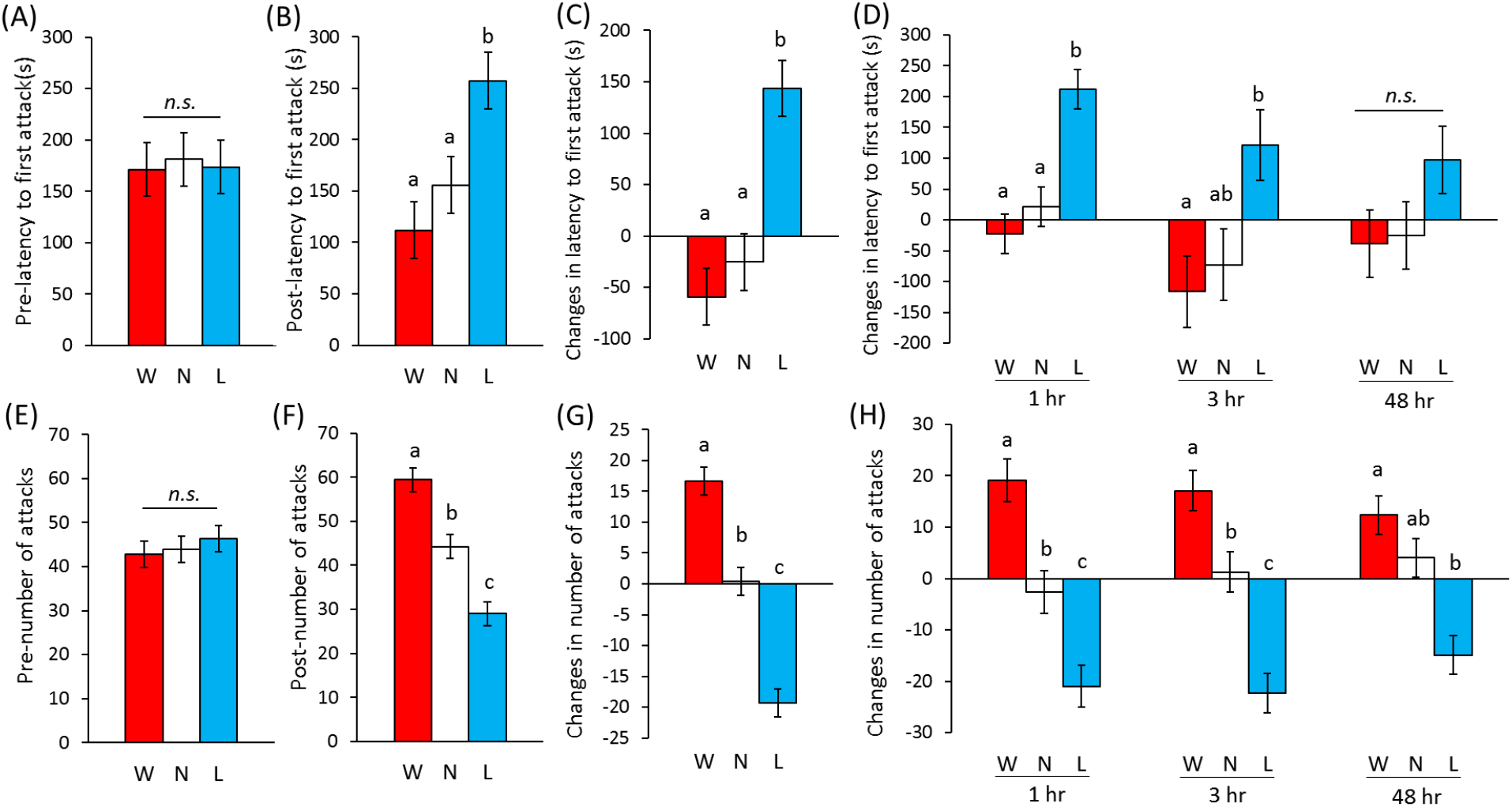
The influence of social experience on aggression. Differences among treatments in aggressive responses toward the non-reversing mirror image: **(A)(E)** before social experience, and **(B)(F)** after social experience. Differences among treatments in how the animals responded to social experience (post-experience behavior minus pre-experience behavior), with respect to **(C)** latency to first attack and **(G)** number of attacks towards the mirror image. Temporal changes in winner-loser effects for **(D)** latency to first attack and **(H)** number of attacks from 1h, 3h to 48h post-experience. Note that ‘latency to first attack’ is inversely related to aggression such that negative changes in latency to first attack indicate increased aggression. Different lowercase letters indicate significant differences between treatments within a given histogram plot (Tukey’s HSD, *P* < 0.05; *n*.*s*. non-significant; W-winners, N-control (no) experience, L-losers; *n* = 75 for each experience).

Relative to their pre-experience behavior, winners, losers and controls attacked the mirror image with higher, lower, and similar frequencies, respectively; all comparisons were significantly different (*P* < 0.001, **Fig. 2G**). Losers also took significantly longer to launch the first attack relative to their pre-experience behavior, a change that was significantly greater than that observed for winners or controls (*P* < 0.001, **Fig. 2C**). While these patterns remained similar across decay time points (**Fig. 2D, Fig. 2H**), there was a significant experience x decay-time interaction for latency to first attack (**Table S1**). Winners’ latency to first attack gradually increased, whereas losers’ latency to first attack gradually decreased between 1h and 48h, suggesting that winner-loser effects were most pronounced at 1h, but slowly decayed after 3h. Also, there were significant differences among lineages in pre-experience aggression, post-experience aggression, and changes in aggressive behavior (***SI Appendix*, Fig. S1A, S1B, Table S1**).

### Social experience and spatial learning

Individuals assigned to each treatment exhibited similar spatial learning performance before social experiences were obtained (**Fig. 3A, 3D, *SI Appendix*, Table S2**). Winners were more likely to pass the spatial learning test than controls at each post-experience time point but not significantly so at 3h (overall: *χ*^*2*^ = 11.14, *P* = 0.002; 1h: *χ*^*2*^= 4.76, *P* = 0.029; 3h: *χ*^*2*^ = 1.64, *P* = 0.205; 48h: *χ*^*2*^ = 6.08, *P* = 0.014, **Fig. 3B, 3C**). Winners also completed the learning task more quickly than losers at each post-experience time point, although they did not perform significantly better at 3 h (losers vs. winners - overall: *P* < 0.001; 1h: *P* = 0.001; 3h: *P* = 0.251; 48h: *P* = 0.039, **Fig. 3E, 3F, *SI Appendix*, Table S2**). Most importantly, winners improved upon their pre-experience performance whereas losers showed virtually no change at all; the difference between winners and losers was significant at all time points except 3 h (losers vs. winners - overall: *P* = 0.001; 1h: *P* = 0.012; 3h: *P* = 0.237; 48h: *P* = 0.009, **Fig. 3G, 3H, *SI Appendix*, Table S2**). There was no significant experience x decay-time interaction on spatial learning, suggesting that performance differences between winners, losers and controls were preserved across time. Lastly, there were significant differences among lineages in pre-experience and post-experience spatial learning performance but not the change in spatial learning performance (***SI Appendix*, Fig. S1C, S1D; Table S2**).

**Figure 3.**
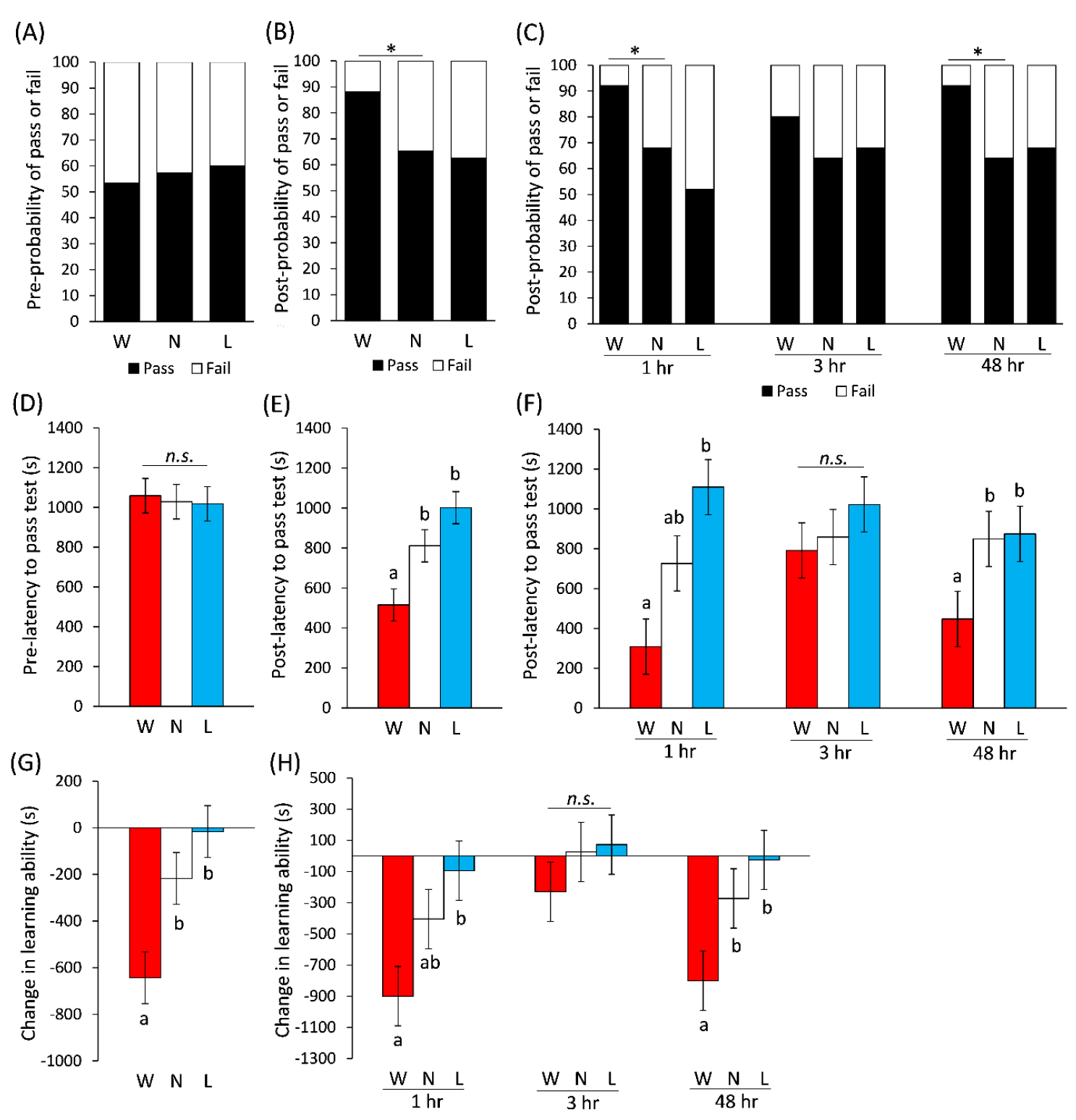
The influence of social experience on spatial learning. Differences among treatments prior to social experience for **(A)** probability of passing/failing the learning test and **(D)** latency (in seconds, *s*) to complete the learning test. Effects of social experience on spatial learning, including **(B)** probability of passing/failing the test, **(E)** latency to complete the learning test and **(G)** change in learning ability (post-experience minus pre-experience performance). Temporal changes in experience effects on spatial learning behavior from 1h, 3h to 48h, including **(C)** probability of passing/failing the test, **(F)** latency to complete the learning test and **(H)** change in learning ability. Note that ‘latency to pass the learning test’ is inversely correlated with learning ability such that negative changes in latency indicate increased learning performance. Asterisk indicates significant difference between treatments (Chi-square test). Different lowercase letters indicate significant differences between treatments within a histogram plot (Tukey’s HSD, *P* < 0.05; *n*.*s*. non-significant; W-winners, N-control (no) experience, L-losers; *n* = 75 for each experience).

### Social experience and risk-avoidance learning

Prior to obtaining social experiences, individuals in the different treatments showed similar performance in the risk-avoidance learning task (**Fig. 4A, 4D, *SI Appendix*, Table S3**). Losers had a higher probability of passing the risk-avoidance learning test than controls at each post-experience time point but not significantly so at 3 h (overall: *χ*^*2*^ = 13.88, *P* < 0.001; 1h: *χ*^*2*^ = 9.14, *P* = 0.003; 3h: *χ*^*2*^ = 2.13, *P* = 0.145; 48h: *χ*^*2*^ = 9.14, *P* = 0.003, **Fig. 4B, 4C**). Losers also solved the learning task faster than winners at all post-experience time points, but the effect was less pronounced at 48 h (losers vs. winners - overall: *P* < 0.001; 1h: *P* = 0.001; 3h: *P* = 0.025; 48h: *P* = 0.174, **Fig. 4E, 4F, *SI Appendix*, Table S3**). Losers improved upon their pre-experience risk learning performance to a greater extent than both winners and controls at each post-experience time point, although the comparison between winners and losers at 48 h was not significant (losers vs. winners - overall: *P* < 0.001; 1h: *P* = 0.001; 3h: *P* = 0.037; 48h: *P* = 0.121; losers vs. controls - overall: *P* = 0.002; 1h: *P* = 0.018; 3h: *P* = 0.037; 48h: *P* = 0.061; **Fig. 4G, 4H, *SI Appendix*, Table S3**). There was no significant experience x decay-time interaction on risk-avoidance learning, suggesting that differences in learning between winners, losers, and controls persisted across post-experience time points. There were significant differences among lineages in pre-experience and post-experience risk avoidance learning performance but not the change in learning performance (***SI Appendix*, Fig. S1E, S1F, Table S3**).

**Figure 4.**
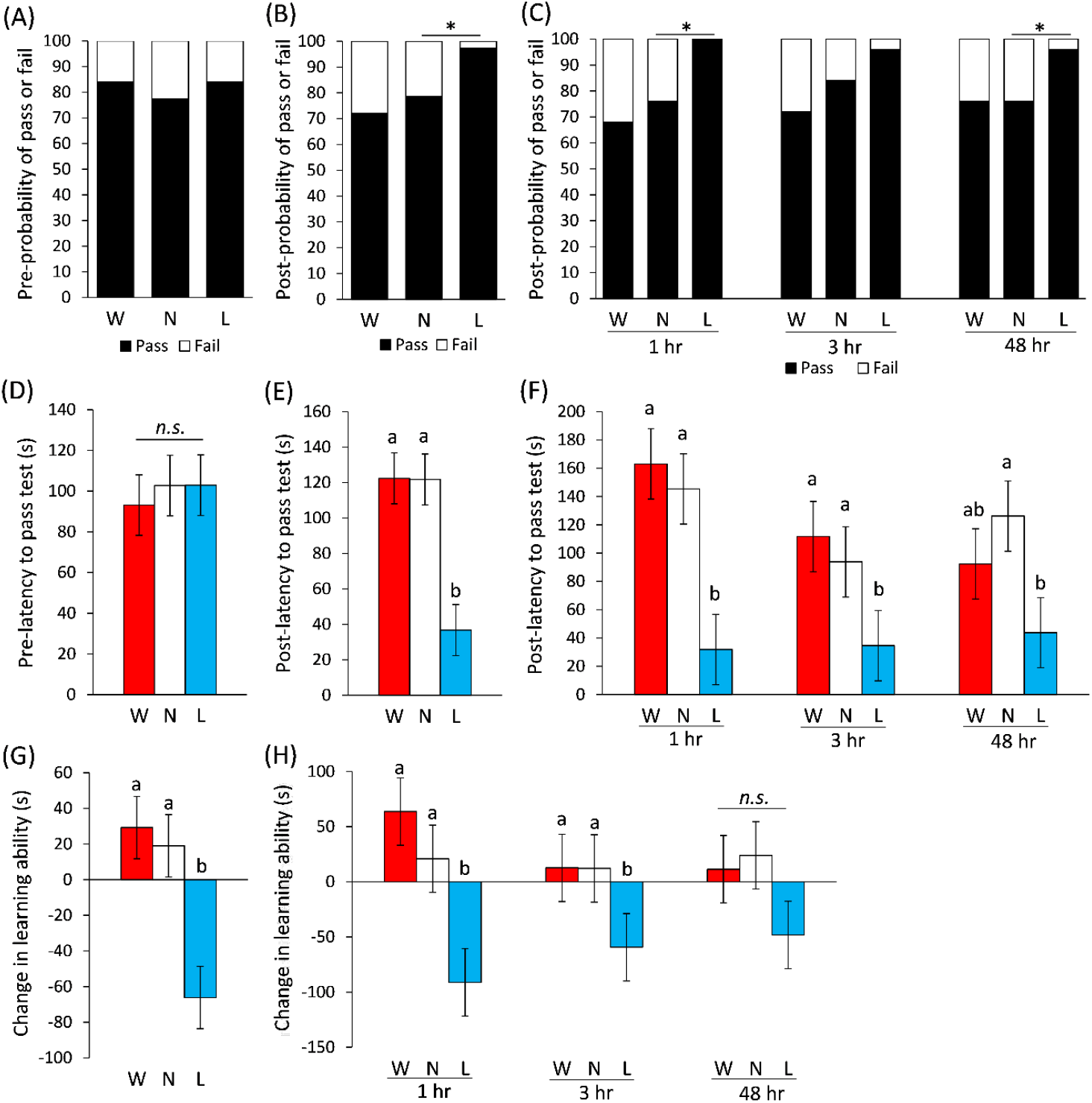
The influence of social experience on risk-avoidance learning. Differences among the treatments prior to social experience for **(A)** probability of passing/failing the learning test and **(D)** latency (in seconds, *s*) to complete the learning test. Effects of social experience on risk-avoidance learning, including **(B)** probability of passing/failing the test, **(E)** latency to complete the learning test, and **(G)** change in learning ability (post-experience minus pre-experience performance). Temporal changes in experience effects on spatial learning behavior from 1h, 3h to 48h, including **(C)** probability of passing/failing the tests, **(F)** latency to complete the learning test and **(H)** change in learning ability. Note that ‘latency to pass learning test’ is inversely correlated with learning ability such that negative changes in latency indicate increased learning performance. Asterisk indicates significant difference between treatments (Chi-square test). Different lowercase letters indicate significant differences between treatments within a histogram plot (Tukey’s HSD, *P* < 0.05; n.s. non-significant; W-winners, N-control (no) experience, L-losers; *n* = 75 for each experience).

### Comparing the effects of social experience on learning abilities

To further investigate how success in the learning tests varied between individuals with different social experiences, we pooled data across decay-time points within the same experience type. We then re-categorized individuals based on learning performance (pass or fail) before and after social experiences (e.g., pass-pass, pass-fail, fail-pass, fail-fail; **Fig. 5**). In the spatial learning test, the pattern of behavioral change in losers was similar to controls (for losers, 26.7% improved [fail to pass] and 24% regressed [pass to fail]; for controls, 26.7% improved and 18.7% regressed). However, winners showed a significantly different pattern than controls (41.3% improved but only 1.3% regressed). Thus, winning dramatically improved spatial learning ability, whereas losing had relatively little effect (**Fig. 5A**).

**Figure 5.**
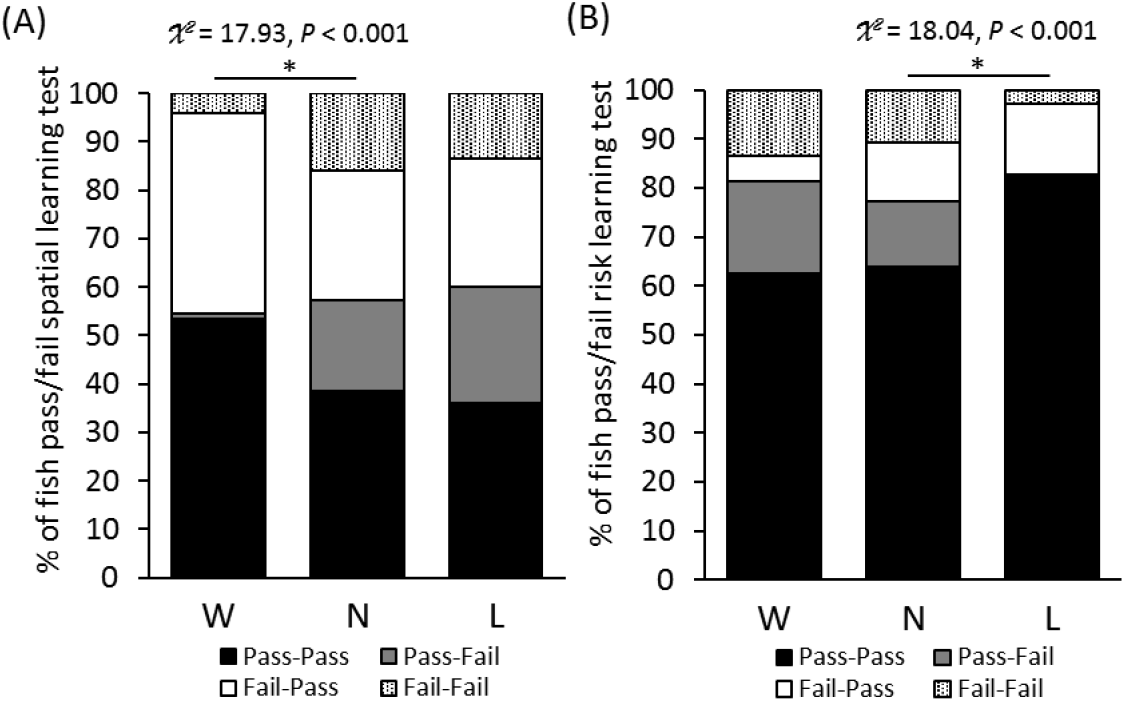
Comparison of probabilities for consecutive pass-fail sequences among the three treatment groups in **(A)** spatial learning ability and **(B)** risk-avoidance learning ability. (Pass-Pass: individuals passed both pre- and post-experience learning tests; Pass-Fail: individuals passed pre-experience learning test but failed post-experience learning test; Fail-Pass: individuals failed pre-experience learning test but passed post-experience learning test; Fail-Fail: individuals failed both pre- and post-experience learning tests; W-winners, N-control (no) experience, L-losers; *n* = 75 for each experience). Asterisk indicates significant difference between treatments (Chi-square test).

For risk-avoidance learning, the pattern of behavioral change in winners was similar to controls (for winners, 5.3% improved and 18.7% regressed; for controls: 13.3% improved and 14.7% regressed). However, 13.3% of losers improved but 0% regressed, which was significantly different from controls. Losing therefore had a strong effect on risk-avoidance learning ability, whereas winning had relatively little effect (**Fig. 5B**).

### Proteomic responses to social experiences

Quantitative proteomics analysis of the forebrain identified 1545 proteins, 23 of which changed significantly in abundance after social experiences were obtained. Of these differentially expressed proteins, four have functions directly related to learning and memory: i) LRRN4 C-terminal-like protein, which is involved in synapse formation, increased 3.9-fold (*P* < 0.001) in losers compared to controls (**Fig. 6**); ii) Ca^2+^/calmodulin-dependent protein kinase type II subunit α, which is involved in long-term potentiation (LTP) and synaptic plasticity, increased 3.5-fold (*P* < 0.001) in winners compared to controls (**Fig. 6**); iii) neuromodulin-like protein, which also participates in LTP, increased 2.5-fold (*P* < 0.001) in losers compared to winners (**Fig. 6**); iv) γ-adducin-like protein, which is involved in LTP and neural firing, increased 2.5-fold (*P* < 0.001) in winners compared to losers (**Fig. 6**). Winners were also found to have a 6-fold higher relative abundance of creatine kinase B-type (CKB) protein, which participates mainly in energy transduction, than controls and losers. In addition to learning and memory, nine differentially expressed proteins were related to cellular processes; five to neural plasticity; two to cell death and apoptosis; two to energy utilization and one to immune function (**Fig. 6**).

**Figure 6.**
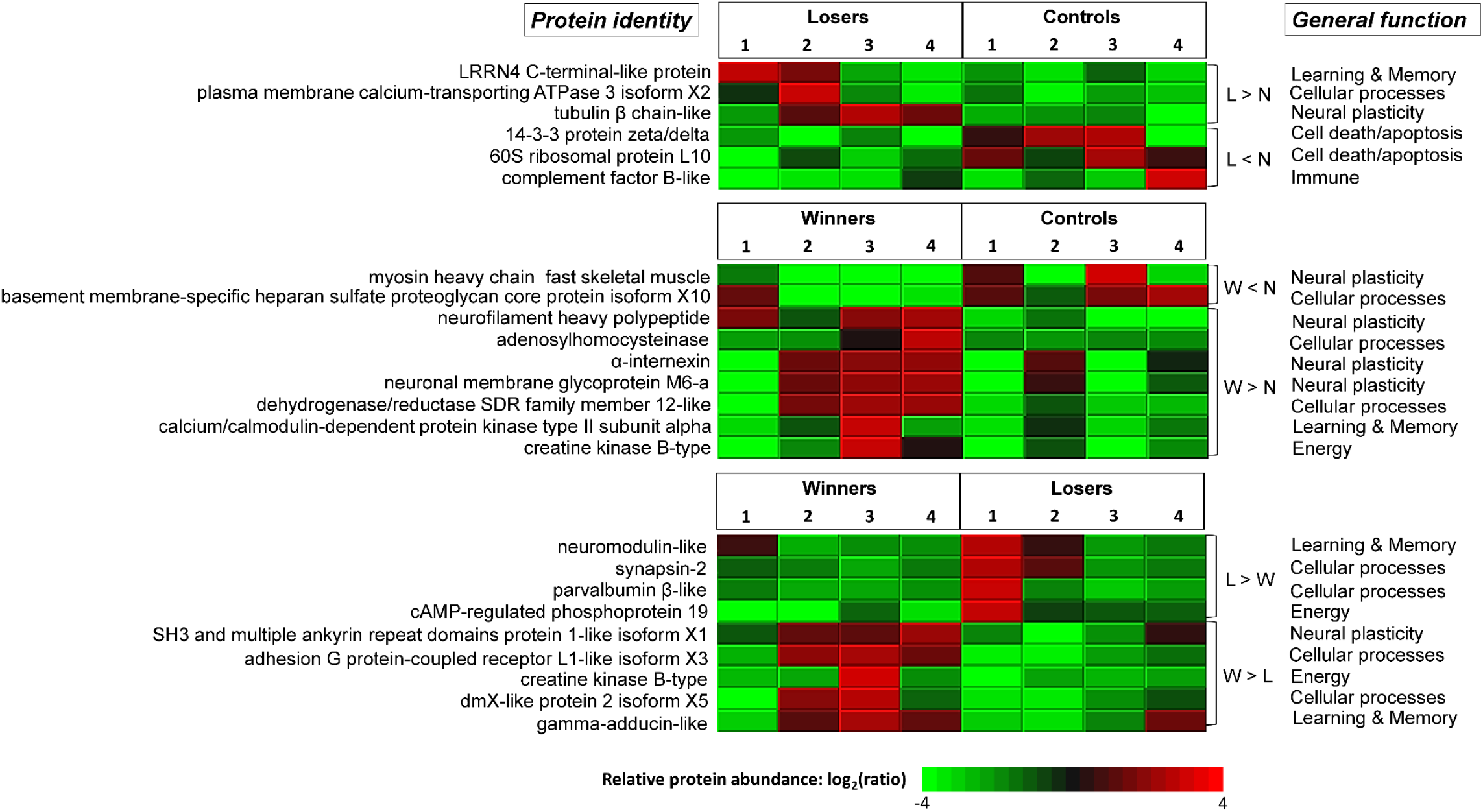
Relative abundances of forebrain proteins that were significantly up- or down-regulated after social experience. Each column represents a single individual (winner *n* = 4, loser *n* = 4, control *n* = 4) and each row represents a unique protein. Red rectangles represent proteins that showed increased expression after social experience and green rectangles represent proteins that showed decreased expression after social experience. Note that the heat map color is based on a log2 scale. (W-winners, N-control (no) experience, L-losers).

Overall, losing affected forebrain expression of proteins that modulate cellular processes (calcium signaling/binding, second messenger production, neurotransmitter release and mobilization of synaptic vesicles), decrease neural plasticity, increase apoptosis and cell death, facilitate recovery from energy deficit (e.g., gluconeogenesis), and mediate learning and memory (e.g., LTP and synapse formation). Winning, however, affected forebrain expression of proteins that mediate cellular processes related to restoration of neuroendocrine homeostasis and signal transduction, increased neuronal plasticity, energy utilization, and learning mechanisms.

## Discussion

We showed that single winning or losing experiences drive markedly different behavioral phenotypes. Predictably, winners increased and losers decreased their aggressive behavior, effects that lasted for at least 48 h. However, we also demonstrated that winning and losing have discernably different effects on learning. Unlike experience-induced changes in aggression, where winners and losers showed opposite responses of similar magnitude, the effects of winning and losing on spatial learning and risk avoidance learning were not symmetrical. Winning increased performance in a spatial learning task but had essentially no effects on performance in a risk-avoidance learning task. On the other hand, losing increased risk-avoidance learning but had no effect on spatial learning. Social experiences also induced pronounced forebrain proteomic responses and, while some proteins were differentially regulated in both winners and losers, many of the proteins were differentially expressed in response to only one type of experience. These data provide evidence that neurobiological responses to winning and losing are not simply ‘opposite sides of the same coin’ but rather, are quite unique. This might thus explain the different (but not necessarily opposite) behavioral phenotypes of winners and losers, especially with respect to learning, and highlights the fact that probing behavioral endpoints other than aggression can illuminate key distinctions in how the brain processes divergent social experiences.

Winner and loser effects were originally defined by changes in aggression after agonistic interactions, and thus most previous research focused on the roles of androgenic hormones and associated receptors in mediating these effects. However, recent studies in invertebrates have revealed that learning and memory may also change in predictable ways with winning and losing experience (21, 31, 32). For instance, fruit flies (*Drosophila melanogaster*) can recognize conspecifics, and losers behave differently when encountering familiar versus unfamiliar opponents, suggesting that learning and memory accompany changes in social status (31). Another study further revealed that repeated losing experiences induced a long-lasting loser effect, which can be blocked by inhibiting protein synthesis, suggesting that the formation of a long-term loser effect requires *de novo* protein synthesis (23). Interestingly, however, repeated winning experience had no such behavioral or physiological consequences in the fruit flies (23). These results imply that different neurobiological mechanisms might govern the expression of winner and loser effects. In crayfish, *Procambarus clarkii*, winner effects can last more than 14 days, and the loser effect can last about 10 days (32). Dominant individuals injected with a 5-HT1 receptor antagonist failed to show the winner effect, whereas subordinate individuals injected with adrenergic-like octopamine receptor antagonist failed to show a loser effect (32). These results provided additional evidence for winner and loser effects being modulated by different neurobiological mechanisms. Furthermore, monoamines, such as serotonin and octopamine, are involved in the expression of various behaviors, ranging from aggression to learning and memory in a diverse array of species, suggesting that winner-loser effects could be mediated by changes in neural processes related to cognition.

Though previous studies have revealed that fighting experience can affect learning and memory (e.g., 19, 20), few studies investigate whether winning and losing experiences influence different types of learning, or the persistence of experience-induced gains and losses in learning ability. Our data revealed that winning and losing not only altered aggression, but also affected spatial and risk-avoidance learning abilities. Temporal patterns of change in aggression and learning also were quite similar, with the effects being most pronounced 1h after fights and gradually decreasing at 3h and 48h. We further discovered that winning and losing independently modulated specific types of learning – spatial learning and risk-avoidance learning, respectively. Spatial learning and memory are mediated primarily by the hippocampus, which provides animals with spatial information of environment and plays important roles in the consolidation of information from short-term memory to long-term memory (33). Risk-avoidance learning is a type of fear conditioning regulated (along with emotional learning) in large part by the amygdala. Both the hippocampus and amygdala also mediate aggressive behavior (34, 35). Most studies hypothesize that winner and loser effects are governed by the same molecule or mechanism, such as testosterone. However, this hypothesis is rarely supported. For instance, Oliveira et al. (4) hypothesized that socially-induced changes in androgen levels should be a causal mediator of both winner and loser effects. They discovered that anti-androgen treatment blocked the winner effect but injecting with androgens failed to rescue the loser effect, suggesting that androgens are involved only in the winner effect. Trannoy et al. (23) showed that winner and loser effects decay over different time courses in fruit flies, which led to the idea that different memory mechanisms might underlie their expression. Our data support this idea and further imply that winning experiences might alter protein expression in the hippocampus and enhance spatial learning, whereas losing experiences might alter protein expression in the amygdala and strengthen risk-avoidance learning. That is, winner and loser effects may not only be governed by different neurobiological mechanisms but might also be mediated by different brain nuclei. Together with previous research, our results suggest that winner-loser effects emerge as a consequence of multiple, perhaps independent neurobiological systems regulating behavioral expression. Because these independent systems can be tuned by the type and intensity of social experience, this is likely to profoundly increase behavioral variation among individuals as experiences accumulate, which can then provide fodder for natural selection. If responses to winning and losing experiences are heritable, as indicated by significant variation among rivulus lineages in experience-induced changes to aggression, this could facilitate the evolution of adaptive neural and behavioral flexibility.

Our proteomics data revealed several promising candidate forebrain proteins associated with social behavior, learning and memory. Identification of brain proteins associated with these behaviors propels dissection of the molecular mechanisms underlying winner and loser effects. For example, creatine kinase B-type (CKB) protein, which is expressed in the hippocampus, cerebellum, and choroid plexus (36), was highly upregulated in winners, but not in losers. CKB mainly participates in energy transduction in the central nervous system but, studies in homozygous knockout mice suggests a critical role for CKB in spatial learning; deficient individuals took longer than wild type individuals to complete the Morris water maze (37). In our study, it is thus possible that winning experiences increase expression CKB in the hippocampus, thereby improving spatial learning abilities. Whether increasing expression of CKB would also alter aggression needs further investigation. Another protein, synapsin-2, which plays a major role in generating synapses and in regulating neurotransmitter release, was significantly upregulated in the forebrains of losers relative to winners. In mice, synapsin-2 is constitutively upregulated in the hippocampus of losers, and its expression is strongly linked with submissive behavior, which is almost uniformly exhibited by animals that experience social defeat (38). Therefore, our results support the idea that losing experiences could upregulate expression of synapsin-2 in the forebrain but, whether expression of synapsin-2 is causally associated with changes in aggression and risk-avoidance learning are unclear.

It is important to note that we only quantified protein abundance 1h after social experience because we observed the most prominent behavioral changes at this time point. Thus, one possible explanation is that *de novo* protein synthesis can cause the observed changes in protein abundance in such short period of time. Alternatively, it is much more likely that specific proteins are rapidly degraded in response to certain signals (39). Studies in rodents revealed that memory consolidation (short-term memory traces being converted to long-term memory) requires not only protein synthesis but also protein degradation (40). In other words, rapid changes in protein abundance could be because certain proteins exist in an unstable state that renders them rapidly degradable in response to a particular stimulus, such as winning or losing experiences.

In summary, we have demonstrated that divergent social experiences alter different learning processes that are mediated by distinct brain nuclei, suggesting that winner effects and loser effects are governed by very different neurobiological mechanisms. We also identified a group of candidate forebrain proteins that might modulate experience-induced changes in behavior. Further experiments that manipulate expression of these candidate proteins in specific brain nuclei (e.g., hippocampus, amygdala and their homologs in other vertebrates) will advance our knowledge about the neural mechanisms underlying experience-induced changes in aggressive behavior and cognitive abilities.

## Materials and Methods

### Study organism

This study used adult hermaphroditic mangrove rivulus, *Kryptolebias marmoratus* (‘rivulus’), from 25 isogenic lineages whose progenitors were wild caught in Belize, the Bahamas, Florida Keys and peninsular Florida. The animals used in this study were two generations removed from field-caught progenitors and were produced via self-fertilization. Individuals were isolated on the day of hatching and kept individually in 1 L translucent plastic containers filled with 750 mL of 25 ppt synthetic seawater (Instant Ocean^®^). Each container was labelled with a unique number for individual identification. Fish were maintained at ambient temperature (27±1°C) on a 12h light: 12h dark photoperiod and fed 2 mL newly hatched brine shrimp (*Artemia*) nauplii every day.

### Providing social experience and quantifying behavior performance

To ensure that individuals received their pre-assigned winning or losing experience, they were fought against much smaller/larger (difference > 2 mm) standard losers/winners that had lost/won several fights against conspecific opponents (random selection procedure, [1]).

In the aggression test, we quantified individuals’ aggressive responses using non-reversing mirror-image stimulation (41, **Fig. 1A**). The latency to initiate aggressive attacks and frequency of aggressive attacks toward mirror image were recorded as aggression indices. In the spatial learning test, individuals were challenged to recall the location of reward (water + brine shrimp nauplii) in a 37 L tank that contained only a layer of moist sponge (1 cm) at the bottom (**Fig. 1B**). This apparatus was modified from Chang et al. (19), and was based on the ecologically relevant premise that rivulus jump or crawl across moist land to seek out water in mangrove forests (42). Fish were given two training sessions (30 min for each session) to become familiar with the environment and to learn the location of the petri dish containing the reward (water + brine shrimp nauplii). During the testing phase, fish were allowed to explore the tank for 30 min to locate the correct petri dish from the previous two training sessions. We considered an individual to have passed or failed the test based on whether it succeeded in locating the correct petri dish; we also recorded the latency to complete the task as a measure of individual spatial learning ability. The risk-avoidance learning test entailed the focal animal being challenged to learn the association between a visual cue (red color, conditioned stimulus [CS]) and an event indicating risk (black corrugated plastic gliding over the tank, unconditioned stimulus [US], **Fig. 1C**). We anticipated that fish would respond to the simulated predator stimulus by seeking shelter and that they would establish an association between red color (CS) and risk signal (US). An individual passed or failed when it sought shelter within 5 min after seeing the red card appear. We also recorded the latency to complete the task as a measure of individual risk-avoidance learning ability. Individuals that failed the spatial learning or risk-avoidance tasks during either training or during the testing phase were also included in the final data set because a failing result was used for comparison between the pre- and post-experience learning performance. Details of each procedure are provided in ***SI Appendix*, Supplementary Material & Methods**.

### Sample preparation and protein quantitation in forebrain

After receiving social experience, rivulus (*n* = 12) were decapitated at 1h in accordance with IACUC standards for euthanasia. Brains were dissected, and forebrains were then separated using a razor blade under a dissecting microscope. Tissues were snap-frozen in liquid nitrogen, stored at −80°C and sent to the University of California, Davis to quantify proteome expression. Protein extraction, protein assays and in-solution trypsin digestion were performed following the protocol established by Kültz and colleagues (43). Detailed procedures regarding sample preparation are provided in ***SI Appendix*, Supplementary Material & Methods**. Protein IDs were mapped to MSMS spectra of particular tryptic peptides using four different search engines, including PEAKS 8.5, Mascot 2.2.7 (Matrix Science, London, UK; version 2.2.07), X!Tandem Alanine (http://www.thegpm.org/tandem/) and Byonic (Protein Metrics, San Carlos, CA, USA; version 2.12). The complete *Kryptolebias marmoratus* proteome (38,516 proteins), an equal number of decoy entries and common contaminants (porcine trypsin, human keratin) were used as the reference database for all searches. Results from all four search engines were consolidated in Scaffold 4.4 (Proteome Software Inc., Portland, OR, USA) and proteins represented by at least 2 unique peptides and meeting a protein level FDR < 1.0% and a peptide level FDR < 0.1% were considered valid IDs. Label-free quantitative profiling of peptide intensities and calculation of relative protein abundances in each sample was performed with PEAKS 8.5. The PEAKS protein quantitation is based on the Top3 approach, which measures the area under the curve of the three most abundant unique peptides for a particular protein from the LC-MS/MS chromatogram. Relative abundance of a peptide in a sample was normalized against the overall abundance of all peptides in that sample.

### Statistical Analysis

General linear models examined whether winners, losers and controls exhibited different levels of aggression before and after social experience, and also whether individuals with different experiences showed variation in the degree to which aggression changed from pre- to post-experience. The response variables were: latency to first attack (ln-transformed to achieve normality) and number of attacks in the pre-experience aggression test or post-experience aggression test, as well as changes in latency to first attack and changes in number of attacks (run in separate models). Type of experience (W, N, L) and decay time (1h, 3h, 48h) were fixed predictor variables. The interaction term, experience x decay time, was also included in the models. Lineage and standard length of focal individuals were included in the model as covariates.

General linear models also examined whether social experience influenced spatial learning and risk-learning abilities. Pre- and post-experience learning behavior, including latency to pass spatial learning and risk learning tasks, and changes in learning abilities were the response variables. Type of experience (W, N, L) and decay time (1h, 3h, 48h) were fixed predictor variables. The interaction term, experience x decay, time was also included in the models. Lineage and standard length of focal individuals were included in the model as covariates.

Tukey’s Honest Significant Difference (HSD) *post hoc* multiple comparisons tests determined differences among levels for the main effects and interactions; results of the Tukey’s HSD tests are presented as *P*-values in parentheses following a description of the differences (**Fig. 2, Fig. 3D-H, Fig. 4D-H**).

Chi-square tests determined whether the probability of successfully passing each learning task differed between winners versus controls and between losers versus controls (**Fig. 3A-C, Fig. 4A-C, Fig. 5**). JMP (v. 12; SAS Institute, Cary, NC, USA) was used for all statistical analyses involving behavior.

Statistical significance for label-free protein quantitation was based on PEAKSQ −log10(*P*-values), which were calculated using a previously developed algorithm that has been optimized for proteomics data (44). In this study, a significance threshold of −log10(*P*-values) ≥ 13 and fold change ≥ 2.0 were applied. All proteomics data, including raw data, metadata, Scaffold file, peptide and protein identifications, and quantitative data are accessible in public proteomics repositories (MassIVE AC: MSV000082806, ProteomeXchange AC: PXD010729). The Scaffold file and zipped PEAKSQ data are accessible via ftp download from MassIVE (AC: MSV000082806).

## Acknowledgements

We thank Caitlin Curtis for assisting in scoring behavior videos. We also thank all members of the Earley laboratory and Kültz laboratory for their comments on the manuscript and their support on this project. This research was supported by Sigma Xi Scientific Research Honors Society (Grants-in-Aid: G201703158955-1084) and College Academy for Research, Scholarship and Creative Activity (CARSCA) committee at the University of Alabama.

