## Supporting Information for "Social experience alters different types of learning abilities controlled by distinct brain nuclei in *Kryptolebias marmoratus*"

### Short title: Social experience and learning ability

Authors: Cheng-Yu Li, Dietmar Kültz, Audrey K. Ward & Ryan L. Earley

Corresponding author:
Cheng-Yu Li


**This PDF file includes:**

Supplementary Material & Methods
Figs. S1 to S2
Tables S1 to S3
Captions for movies S1 to S5
Captions for databases S1
References for SI reference citations

**Other supplementary materials for this manuscript include the following:**

Movies S1 to S5
Datasets S1

**Supplementary Material & Methods**

***Study organism***

This study used adult hermaphroditic mongrove rivulus, *Kryptolebias marmoratus*, of 25 isogenic lineages (BCV45-52, BP11, BP21, BUN1, BUN5, BWS11, CF1-1, CF15, CRWL19, DC9, ERIN11, FDS1-1, FDS3-E16, HAM2, IRP5, LMC3, MES11, NEL10, NNKN1, NUKE7, OSR7-8, RHL12, RHL5, VERO1 and WEED3), from Belize, the Bahamas, Florida Keys and peninsular Florida. Individuals were isolated on the day of hatching and kept individually in 1000mL translucent plastic containers (maintenance container) filled with 750mL of 25 ppt synthetic seawater (Instant Ocean®). Each container was labelled with a unique number for individual identification. Fish were maintained at ambient temperature (27±1°C) on a 12h light: 12h dark photoperiod and fed 2mL newly hatched brine shrimp (*Artemia*) nauplii every day. The average (± SEM) age of experimental animals was 324.85 ± 21.14 days; the average (± SEM) standard length of experimental animals was 24.48 ± 0.07 mm.

***Experimental design***

To investigate the temporal influences of social experience on behavioral variation in aggression and learning-memory, we conducted a full factorial experiment with 3 experience levels and 3 decay-time levels. Fish ($n$ = 625) were allocated to three experience treatments (W: social victory; L: social defeat; N: control experience). Individuals were subjected to behavioral tests before (on Day 1) and after social experience (1h, 3h and 48h on Day 15-17, according to pre-assigned decay-time treatments). Animals were divided evenly among three groups, which were exposed to the aggression, spatial learning or risk-aversion learning tests. To ensure that focal animals obtain winning or losing experience, we fought them against much smaller or larger (Δ > 2 mm) opponents (random selection, 1). Quantifying behaviors before and after social competition provides information about 'baseline' individual differences in behavior and experience-induced changes in aggression and learning abilities. Quantifying behaviors at 1h, 3h and 48h after a fight was used to investigate temporal changes in behavior following winning or losing experience. Instead of quantifying behavior immediately after social experience, we allowed fish to rest for one hour to recover from an aggressive interaction, which was meant to minimize the influence of fatigue on the animals' motivation to engage with a conspecific or the learning tasks. A total of 12 individuals (winner, $n = 4$; losers, $n = 4$, controls [no experience], $n = 4$) were used in proteomic analysis. We quantified forebrain proteome expression 1h after a winning or losing experience because fish exhibited the most pronounced behavioral changes in aggression and learning at 1h post-experience.

***Providing social experience***

To ensure that individuals won or lost as pre-assigned, they were fought against much larger/smaller (difference > 2 mm) standard winners/losers that had won/lost several fights against conspecific opponents. Before experience training, fish were transferred from the maintenance container to one of two symmetrical compartments (randomly selected) of a $12 \times 8 \times 20$ cm$^3$ contest aquarium (water 13 cm deep and 2 cm of gravel) separated by an opaque partition from a standard winner or standard loser in the other compartment. All fish were given 30 min to acclimate to the environment. After removing the opaque partition, a loss was completed when the experimental individual retreated from the standard winner's display/attack and quickly swam away; while a win was completed when the standard loser retreated from the experimental individual's display/attack and quickly swam away. All fish were returned to their maintenance containers after experience training. Control individuals were treated in exactly the same way as the others, except with no opponent in the contest aquarium, so that they received the same amount of handling as the other experimental individuals.

***Aggression test***

We quantified the latency to initiate aggressive attacks and frequency of aggressive attacks using a non-reversing mirror-image stimulation. All social tests were conducted in a standard aquarium ($12 \times 12 \times 12$ cm$^3$, containing water 11 cm deep and 0.5 cm of gravel). Aggressive behavior was quantified by placing a non-reversing mirror in a standard aquarium (2, 3). A non-reversing mirror was made by gluing two first-surface mirrors (4.5 × 8.0 $cm^2$) at their edges at a 90-degree angle and then attached to the corner of the standard aquarium. A transparent glass divider (6.5 × 8.0 $cm^2$) was placed between the experimental fish and the non-reversing mirror (**Fig. 1A**) to prevent individuals from seeing multiple images during the aggression test. Individuals were allowed to acclimate for 30 min behind an opaque partition. After the acclimation period, the partition was removed, and the fish was allowed to interact with the non-reversing mirror image. Behaviors were recorded by camcorders (DV CR303, Samsung, Seoul, South Korea). The latency to initiate aggressive attacks and frequency of aggressive attacks were recorded as aggression indices.

***Spatial learning test***

Individuals were challenged to recall the location of water in a T-maze (**Fig. 1B**), which was modified from Chang et al. (4); this modified T-maze focused on the ecologically relevant premise that rivulus jump or crawl across moist land to seek out water in mangrove forests (5). The maze consisted of two petri dishes 9.0 × 1.0 $cm^2$ symmetrically arranged at the corner of a 40 × 20 × 26 $cm^3$ tank lined with 1.0 cm of white sponge in order to maintain body moisture during training and testing procedures. Above each petri dish was a cutout shape (triangle or square) made of black corrugated plastic to provide a visual cue to associate with the water resource. The visual cues and resource dish were randomly switched between each testing individual to prevent experimental bias and to negate the possibility that individuals were succeeding because of a natural tendency to jump in one particular direction. In the center of the tank was a square piece of black corrugated plastic (9.0 × 9.0 $cm^2$) onto which the fish was transferred prior to the beginning of each session. Fish were transferred in an opaque petri dish with black plastic cover to minimize stress. Fish were then allowed to acclimate for 1 min, after which time the cover was removed and the testing fish could begin exploring and navigating toward the water resource.

During the training phase, individuals needed to locate, within 30 min, the one petri dish that was filled with water and a few drops of brine shrimp nauplii; it is not unusual for the fish to voluntarily stay out of water for this amount of time, and they can live out of water in a moist environment for at least 2 months (6). If fish failed to locate the dish containing water during the 30 min period, the training session was stopped, and this individual was forced to jump into the correct dish by probing it gently with a transfer pipette. Once the fish jumped into the water-filled dish, it was allowed to rest for 15 min to become familiar and create associations with the visual cues in the environment. Following each training period, individuals were returned back to the maintenance container for 1h rest. Fish were given two training sessions to become familiar with the environment and to learn the location of the reward (water + brine shrimp nauplii). The testing phase was designed to determine whether an individual was capable of learning the location of water associated with a specific visual cue. During this phase, neither petri dish was filled with water or brine shrimp, and fish were allowed to explore the tank for 30 min to locate the correct petri dish from the previous two training sessions. The visual cues were placed in the same spot as the individual experienced during the training phase for helping fish to locate the correct petri dish. We considered an individual to have passed or failed the test based on whether it succeeded in locating the correct petri dish; we also recorded the latency to complete the task as a variable to quantify individual spatial learning ability. Behaviors were recorded by camcorders (DV CR303).

***Risk-avoidance learning test***

The risk-avoidance learning tests entailed the focal animal being challenged to learn the association between a visual cue (red color, conditioned stimulus [CS]) and an event indicating risk (a black corrugated plastic gliding over the tank, unconditioned stimulus [US], **Fig. 1C**). This test was conducted in a standard tank ($12.0 \times 8.0 \times 20.0$ cm$^3$) filled with room-heated, 25 ppt water and 5 cm of gravel. At one end of tank there was a shelter made by black corrugated plastic and at the other end of the tank there was a window ($4.0 \times 8.0$ cm$^2$) for fish to observe the red color. Before the training phase, individuals were allowed to acclimate and explore the environment for 30 min until they were freely swimming in the tank. During the training phase, a red card appeared in the window along with a black corrugated plastic (30.0 × 10.0 $cm^2$) gliding over the tank four times, which was meant to simulate a predator approaching from above. Individuals were given two sessions of CS-US pairing to learn the association. We anticipated that fish would respond to the simulated predator stimulus by either freezing or seeking shelter and that they would establish an association between red color (CS) and risk signal (US). In the testing phase, fish were exposed only to the red color (CS); if fish established the association between CS-US, we expected that they would exhibit freezing behavior or seek shelter. We considered an individual to have passed or failed the test based on whether it exhibited freezing behavior or sought shelter within 5 min after seeing the red card appear in the window. We also recorded the latency to complete the task to quantify individual risk-avoidance learning ability. Behaviors were recorded by camcorders (DV CR303).

Individuals that failed the spatial learning or risk-avoidance tasks during either training or during the testing phase were also included in final data, as a failing result was still used for comparison between the pre- and post-experience learning ability.

***Forebrain protein sample preparation***

After receiving social experience, mangrove rivulus ($n$ = 12) were decapitated at 1h in accordance with IACUC standards for euthanasia. Brains were micro-dissected, and forebrains were then separated using a razor blade under a dissecting microscope. Tissues were snap-frozen in liquid nitrogen and stored at -80°C and sent to the University of California, Davis, to quantify proteome expression. Protein extraction, protein assays and in-solution trypsin digestion were performed following the protocol as reported previously (7). Two hundred nanograms of tryptic peptides from each sample were injected with a nanoAcquity sample manager (Waters, Milford, MA, USA) and trapped for 1 min at 15 µl/min on a Symmetry trap column (Waters, MA, USA). They were then separated on a BEH C18 column (1.7 µm particle size, 250 mm×75 µm, Waters, MA, USA) using a 125-min linear gradient ranging from 3% to 35% acetonitrile by reversed phase liquid chromatography using a nanoAcquity UPLC (Waters, MA, USA). Nano-electrospray ionization (nESI) was achieved by elution from a New Objective Pico Tip emitter (FS360–20-10-D-20, Woburn, MA, USA) into a nanoESI source fitted on an ImpactHD UHR-QTOF mass spectrometer (Bruker Daltonics, Bremen, Germany). Batch-processing of samples was controlled with Hystar 4.1 software (Bruker Daltonics).

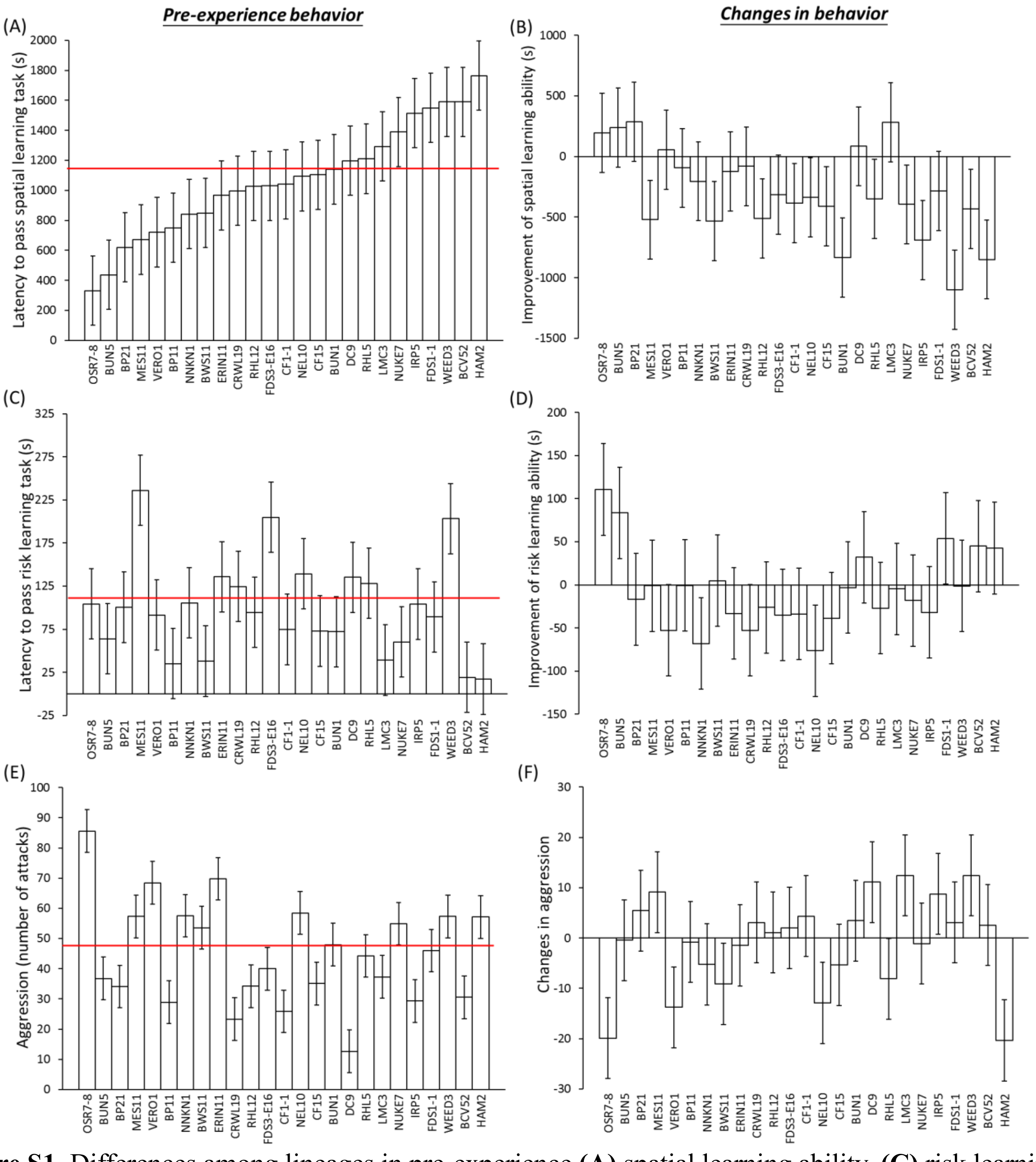


**Figure S1.** Differences among lineages in pre-experience **(A)** spatial learning ability, **(C)** risk learning ability and **(E)** aggression. Differences among lineages in changes of behavior, including **(B)** spatial learning ability, **(D)** risk-avoidance learning and **(F)** aggression. Red lines represent the average behavioral performance across all lineages. Note that "latency to pass learning test" is inversely correlated with learning ability such that negative changes in latency represent increased learning performance.

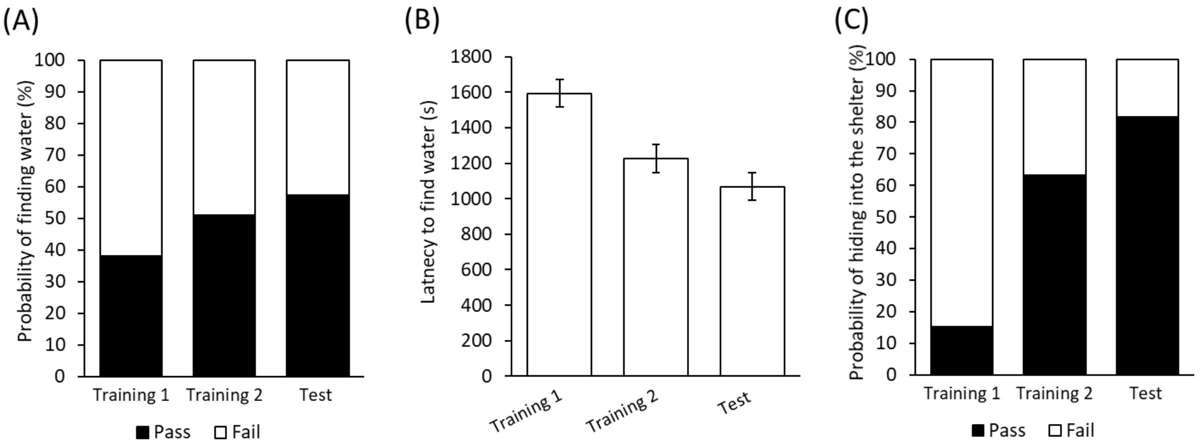


**Figure S2.** Overall learning performance in spatial learning test and in risk-avoidance learning test between training sessions and testing session. Individuals that received two training sessions had significant improvements in the probability of finding water (**A**) and in the latency to find water (**B**). Individuals that received two training sessions also had significant improvements in learning the association between a visual cue (red color, CS) and an event indicating risk (a black corrugated plastic gliding over the tank, US) (**C**). Note that "latency to find" is inversely correlated with learning ability such that negative changes in latency represent increased learning performance.

**Table S1.** Models examining differences of social experiences on individuals' aggressive behavior and changes in aggressive behavior.

| Variable | *df* | *b*±*SE* | *F* | *P* |
|---|---|---|---|---|
| **Pre-experience latency to first attack** | | | | |
| Experience type (W, N, L) | 2, 191 | | 1.74 | 0.178 |
| Decay time type (1h, 3h, 48h) | 2, 191 | | 2.13 | 0.122 |
| Experience x Decay time | 4, 191 | | 0.92 | 0.452 |
| Lineage | 24, 191 | | **2.56** | **<0.001*** |
| Standard length | 1, 191 | 0.02±0.05 | 0.19 | 0.661 |
| **Pre-experience total attack numbers** | | | | |
| Experience type (W, N, L) | 2, 191 | | 1.44 | 0.240 |
| Decay time type (1h, 3h, 48h) | 2, 191 | | 1.24 | 0.290 |
| Experience x Decay time | 4, 191 | | 0.30 | 0.876 |
| Lineage | 24, 191 | | **5.26** | **<0.001*** |
| Standard length | 1, 191 | 1.34±1.15 | 1.35 | 0.246 |
| **Post-experience latency to first attack** | | | | |
| Experience type (W, N, L) | 2, 191 | | **25.52** | **<0.001*** |
| Decay time type (1h, 3h, 48h) | 2, 191 | | **3.06** | **0.049*** |
| Experience x Decay time | 4, 191 | | **4.00** | **0.004*** |
| Lineage | 24, 191 | | **3.05** | **<0.001*** |
| Standard length | 1, 191 | -0.06±0.04 | 2.10 | 0.149 |
| **Post-experience total attack numbers** | | | | |
| Experience type (W, N, L) | 2, 191 | | **41.95** | **<0.001*** |
| Decay time type (1h, 3h, 48h) | 2, 191 | | 0.58 | 0.559 |
| Experience x Decay time | 4, 191 | | 0.78 | 0.540 |
| Lineage | 24, 191 | | **3.79** | **<0.001*** |
| Standard length | 1, 191 | 0.62±1.10 | 0.31 | 0.576 |
| **Changes in latency to first attack** | | | | |
| Experience type (W, N, L) | 2, 191 | | **15.96** | **<0.001*** |
| Decay time type (1h, 3h, 48h) | 2, 191 | | 3.01 | 0.052 |
| Experience x Decay time | 4, 191 | | 0.40 | 0.811 |
| Lineage | 24, 191 | | **1.94** | **0.008*** |
| Standard length | 1, 191 | -1.83±12.76 | 0.02 | 0.886 |
| **Changes in total attack numbers** | | | | |
| Experience type (W, N, L) | 2, 191 | | **71.12** | **<0.001*** |
| Decay time type (1h, 3h, 48h) | 2, 191 | | 0.19 | 0.825 |
| Experience x Decay time | 4, 191 | | 0.96 | 0.432 |
| Lineage | 24, 191 | | **2.17** | **0.002*** |
| Standard length | 1, 191 | -0.72±1.00 | 0.51 | 0.475 |

Standard length and lineages were included in models as covariates, Experience type: W-winning experience; N-no experience; L-losing experience. Changes in behavior: (post-experience behavior) – (pre-experience behavior) (*df*: the degree of freedom; * $P < 0.05$).

**Table S2.** Models examining differences of social experiences on individuals' spatial learning behavior and changes in learning performance.

| Variable | *df* | *b±SE* | *F* | *P* |
|---|---|---|---|---|
| **Pre-experience** | | | | |
| **latency to pass spatial learning task** | | | | |
| Experience type (W, N, L) | 2, 191 | | 0.92 | 0.399 |
| Decay time type (1h, 3h, 48h) | 2, 191 | | 2.19 | 0.114 |
| Experience x Decay time | 4, 191 | | 0.61 | 0.656 |
| Lineage | 24, 191 | | **2.57** | **<0.001*** |
| Standard length | 1, 191 | 11.21±37.42 | 0.09 | 0.765 |
| **Post-experience** | | | | |
| **latency to pass spatial learning task** | | | | |
| Experience type (W, N, L) | 2, 191 | | **10.41** | **<0.001*** |
| Decay time type (1h, 3h, 48h) | 2, 191 | | 1.66 | 0.193 |
| Experience x Decay time | 4, 191 | | 1.80 | 0.130 |
| Lineage | 24, 191 | | **1.93** | **0.008*** |
| Standard length | 1, 191 | -32.98±35.00 | 0.89 | 0.347 |
| **Changes in spatial learning abilities** | | | | |
| Experience type (W, N, L) | 2, 191 | | **8.54** | **<0.001*** |
| Decay time type (1h, 3h, 48h) | 2, 191 | | **3.84** | **0.023*** |
| Experience x Decay time | 4, 191 | | 0.62 | 0.649 |
| Lineage | 24, 191 | | 1.22 | 0.232 |
| Standard length | 1, 191 | -44.19±50.43 | 0.77 | 0.382 |

Standard length and lineages were included in models as covariates, Experience type: W-winning experience; N-no experience; L-losing experience. Changes in behavior: (post-experience behavior) – (pre-experience behavior). *df*: the degree of freedom; * $P < 0.05$.

**Table S3.** Models examining differences of social experiences on individuals' risk-avoidance learning behavior and changes in learning performance. (ESM)

| Variable | *df* | *b*±*SE* | *F* | *P* |
|---|---|---|---|---|
| **Pre-experience** | | | | |
| **latency to pass risk-avoidance learning task** | | | | |
| Experience type (W, N, L) | 2, 191 | | 0.02 | 0.980 |
| Decay time type (1h, 3h, 48h) | 2, 191 | | 1.16 | 0.315 |
| Experience x Decay time | 4, 191 | | 0.25 | 0.912 |
| Lineage | 24, 191 | | **2.11** | **0.003*** |
| Standard length | 1, 191 | -14.57±5.72 | **6.50** | **0.012*** |
| **Post-experience** | | | | |
| **latency to pass risk-avoidance learning task** | | | | |
| Experience type (W, N, L) | 2, 191 | | **14.25** | **<0.001*** |
| Decay time type (1h, 3h, 48h) | 2, 191 | | 1.83 | 0.163 |
| Experience x Decay time | 4, 191 | | 1.10 | 0.356 |
| Lineage | 24, 191 | | **2.71** | **<0.001*** |
| Standard length | 1, 191 | -6.16±5.33 | 1.33 | 0.250 |
| **Changes in risk-avoidance learning abilities** | | | | |
| Experience type (W, N, L) | 2, 191 | | **8.39** | **<0.001*** |
| Decay time type (1h, 3h, 48h) | 2, 191 | | 0.10 | 0.903 |
| Experience x Decay time | 4, 191 | | 0.70 | 0.595 |
| Lineage | 24, 191 | | 0.84 | 0.686 |
| Standard length | 1, 191 | 8.42±7.22 | 1.36 | 0.245 |

Standard length and lineages were included in models as covariates, Experience type: W-winning experience; N-no experience; L-losing experience. Changes in behavior: (post-experience behavior) – (pre-experience behavior). *df*: the degree of freedom; * $P < 0.05$.

**Movie S1.** An example of focal fish's (CY#581) performance in the first training of spatial learning task. This individual found water/food reward at 11 minutes 10 seconds.

**Movie S2.** An example of the same focal fish's (CY#581) performance in the second training of the spatial learning task. This individual only spent 1 minute 51 seconds to find water/food reward.

**Movie S3.** The same focal fish's (CY#581) performance in the test session of the spatial learning task. This individual passed the test at 1 minute 55 seconds (i.e., jumped on to the correct petri-dish).

**Movie S4**. An example of focal fish's performance in the first training of risk-avoidance learning task. At 24 seconds, this individual hid into the shelter after observing the conditioned stimulus ([CS], red color appeared on the left window) and unconditioned stimulus ([US], black corrugated plastic gliding over the tank to simulate a risk event) appeared in order.

**Movie S5**. An example of focal fish's performance in the test session of risk-avoidance learning task. At 27 seconds, this individual hid into the shelter after observing the conditioned stimulus ([CS], red color appeared on the right window).

**Database S1**. Behavior data from aggression test, spatial learning test and risk-avoidance learning test are listed in this excel spreadsheet.
